# Droplet bioprinting of acellular and cell-laden structures at high-resolutions

**DOI:** 10.1101/2023.11.18.567660

**Authors:** Puskal Kunwar, Ujjwal Aryal, Arun Poudel, Daniel Fougnier, Zachary J Geffert, Rui Xie, Zhen Li, Pranav Soman

**Affiliations:** Syracuse University, Biomedical, and Chemical Engineering Department, Syracuse, New York, 13210, USA; BioInspired Institute, Syracuse, New York, 13210, USA; Clemson University, Department of Mechanical Engineering, Clemson, SC 29634, USA

**Keywords:** Digital light processing, bioprinting, hydrogel, cell encapsulation, cell seeding, high resolution, low volume, no waste

## Abstract

Advances in Digital Light Processing (DLP) based (bio) printers have made printing of intricate structures at high resolution possible using a wide range of photosensitive bioinks. A typical setup of a DLP bioprinter includes a vat or reservoir filled with liquid bioink, which presents challenges in terms of cost associated with bioink synthesis, high waste, and gravity-induced cell settling, contaminations, or variation in bioink viscosity during the printing process. Here, we report a vat-free, low-volume, waste-free droplet bioprinting method capable of rapidly printing 3D soft structures at high resolution using model bioinks. A multiphase many-body dissipative particle dynamics (mDPD) model was developed to simulate the dynamic process of droplet-based DLP printing and elucidate the roles of surface wettability and bioink viscosity. Process variables such as light intensity, photo-initiator concentration, and bioink formulations were optimized to print 3D soft structures (∼0.4 to 3 kPa) with an XY resolution of 38 ± 1.5 μm and Z resolution of 237±5.4 μm. To demonstrate its versatility, droplet bioprinting was used to print a range of acellular 3D structures such as a lattice cube, a Mayan pyramid, a heart-shaped structure, and a microfluidic chip with endothelialized channels. Droplet bioprinting, performed using model C3H/10T1/2 cells, exhibited high viability (90%) and cell spreading. Additionally, microfluidic devices with internal channel network lined with endothelial cells showed robust monolayer formation while osteoblast-laden constructs showed mineral deposition upon osteogenic induction. Overall, droplet bioprinting could be a low-cost, no-waste, easy-to-use, method to make customized bioprinted constructs for a range of biomedical applications.

## Introduction

3D bioprinting encompasses a set of technologies that have been widely used to generate tissue constructs for applications in tissue engineering, regenerative medicine, drug screening, and disease modeling.^[1–7]^ Among the many bioprinting methods, digital light projection (DLP) based bioprinting methods are capable of printing both acellular and cell-laden 3D architecture at high resolutions, speeds, and overall fidelity.^[3,4,7–10]^ These methods use a liquid crystal display or a digital micromirror device (DMD) to generate digital masks that can spatially pattern and project light onto a vat or reservoir filled with liquid photo-sensitive bioink to print 3D constructs via rapid crosslinking in a layer-by-layer manner.^[11–13]^ Depending on the optical setups and bioink formulations, structures with a high XY resolution of 50-100 μm are possible. Vat-based DLP bioprinting has already been used to develop 3D tissue models, ^[12,14]^ hydrogel-based microfluidic chips, heart valves, and scaffolds for joint and ligament regeneration, among other constructs.^[15]^ However, many challenges arise due to the use of vat in DLP bioprinting. Typically, the entire vat must be filled with expensive photo-sensitive bioinks, and after the printing process is finished, the remaining bioinks are discarded. Unlike printing of commercial resins, bioinks are often synthesized in-house and can incur high costs based on the types of formulations and cell types, so every effort should be made to minimize waste. Even for short printing times, cells within bioinks undergo gravity-induced settling which can result in uneven cell distributions within the printed structures. Longer print durations increase the likelihood of (i) contamination and cell death, (ii) low cell viability due to longer exposure to photo-initiators, (iii) undesired bioink gelation due to changes in viscosity; all these could lead to failed prints.^[16][17]^ With vat-bioprinting, screening and/or optimization of bioinks for specific applications also remains expensive and wasteful; as a result, only limited variables are typically tested. To address these challenges, we report a vat-free, low-volume droplet bioprinting method capable of rapidly printing acellular and cell-laden 3D soft structures at high resolution using model bioinks such as poly(ethylene glycol)-diacrylate (PEGDA) and gelatin methacrylate (GelMA).

## Results and Discussion

### 1. Strategy, Setup Design, and Optimization

**Figure 1A** illustrates the schematic of DLP-based droplet bioprinting. A 405 nm light, spatially patterned using a Digital Micromirror Device (DMD), is passed through a light-transparent and oxygen-permeable PDMS window to enable crosslinking of a single layer followed by synchronized movement of the L-shaped stage. The first step in printing a 3D construct is the generation of a CAD model (SolidWorks), then MATLAB code is employed to slice the 3D structure into 2D PNG images that serve as digital masks; 1-bit images to define the spatial distribution for corresponding 2D slices. These processed images are supplied to a custom LabVIEW code which precisely coordinates stage movements and DMD-generated light patterns to print the 3D construct in a layer-by-layer manner. The oxygen permeability of the PDMS window inhibits crosslinking at the interface and generates a ‘dead zone’ to facilitate continuous printing. Before printing, a bioink droplet (photosensitive bioink), whose volume is defined by the CAD model, is placed in the fabrication window – the space between the PDMS and stage; this forms a three-phase contact line at the interface between the bioink, PDMS, and air. The surface energy of the PDMS acts along the solid surface, while the interfacial energy between the PDMS and the bioink acts in the opposite direction. Tangentially, the surface tension of the polymer acts on the drop surface. The resulting vector force causes the bioink to form a dome-like shape due to these combined forces, which are characterized by a contact angle θ_1_, which is the angle formed at the intersection of a bioink, air, and PDMS membrane at the three-phase boundary (**Figure 1B**). Upward stage movement draws the bioink towards the fabrication window, then the stage moves down to achieve the desired layer thickness before light irradiation and photo-crosslinking. As the printing process progresses, another three-phase contact line is formed between the freshly crosslinked structure, bioink, and air. This new contact line defines a different contact angle represented by θ_2_(**Figure 1B**). The presence of these two three-phase contact lines results in the formation of a meniscus surrounding the freshly crosslinked structure (**Video V1**). During printing, the bioink from the meniscus is consumed and the method allows adding fresh ink at any point during the fabrication process. In this work, we choose an up-down motion instead of a continuous upward motion of the stage, to help draw the liquid bioink into the fabrication window **(Figure 1C)**. This process is repeated multiple times to print the final structure in a layer-by-layer fashion.

**Figure 1.**
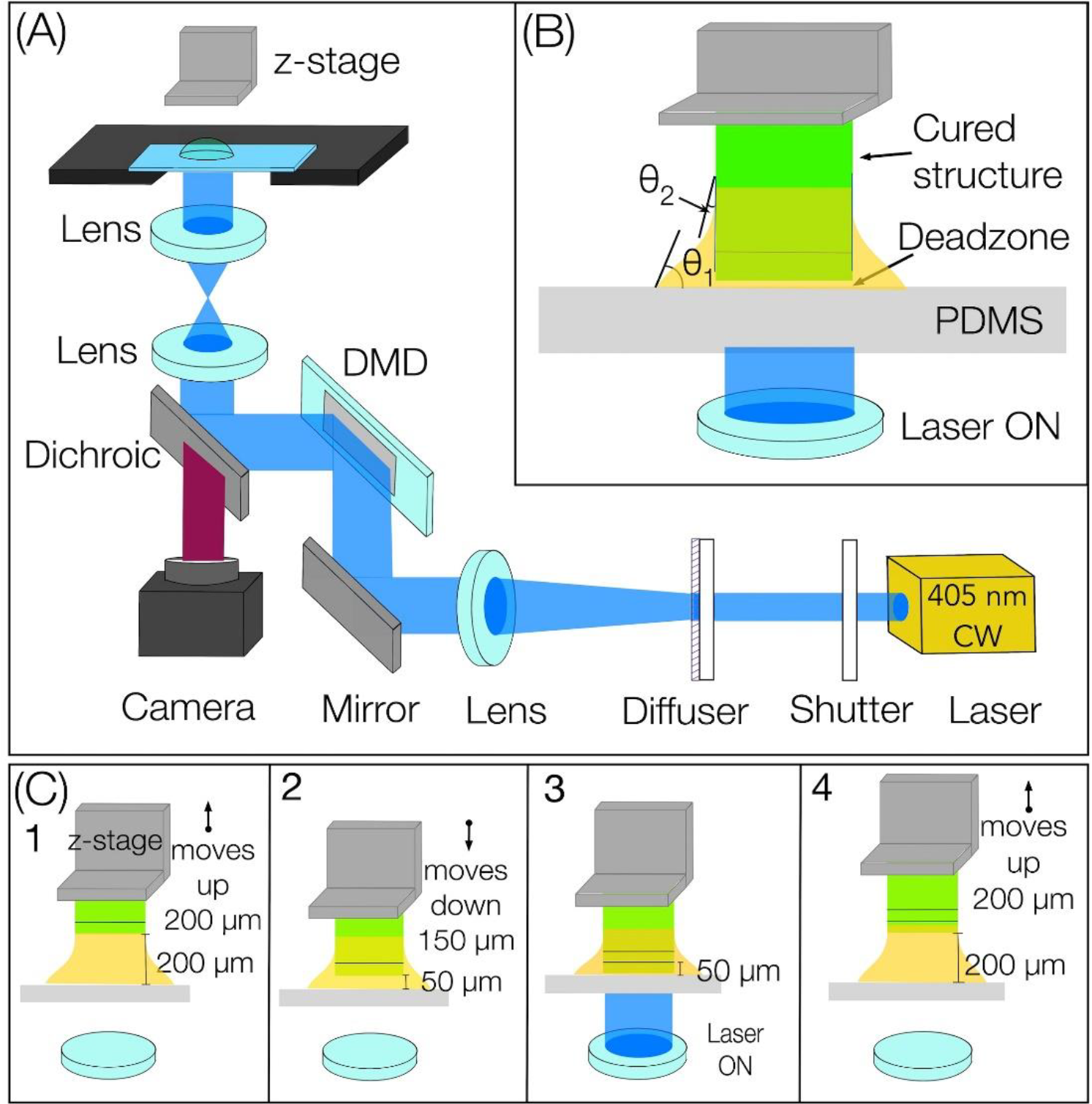
**(A)** The schematic demonstrates the vat-free DLP-based droplet bioprinting **(B)** The figure showcases the meniscus, which is governed by two contact angles: θ_1_ represents the contact angle formed by the bioink on the PDMS surface, while θ_2_ represents the contact angle formed by the bioink on the crosslinked structure **(C)** Inset 1-4 shows the upward and downward movement of the stage facilitating the drawing of bioink in the fabrication area.

For a typical structure with 100 layers, the volume of excess bioink left behind is less than that required to print 1 layer is left behind (waste ∼< 1% of initial volume).

### 2. Simulation studies of the droplet printing process

Designing experimental configurations for achieving the seamless and uninterrupted 3D printing of diverse bioinks can prove to be both costly and time intensive. Since viscosity, surface forces between bioink, printhead, and PDMS and stage speed, all contribute towards high print fidelity, we developed a multiphase many-body dissipative particle dynamics (mDPD) model ^[18,19]^ to simulate the dynamic process of droplet bioprinting with a system setup shown in **Figure 2A(i)**.

**Figure 2.**
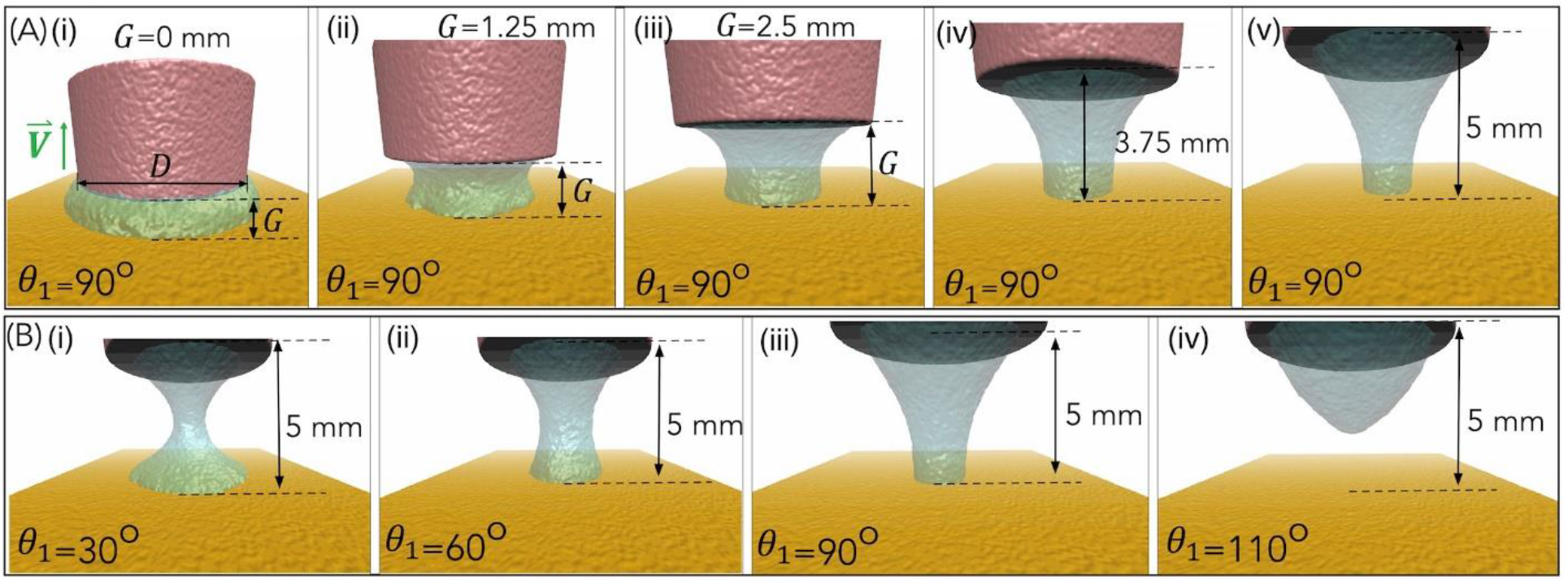
**(A**). A sequence of simulation snapshots depicting the movement of liquid into the fabrication area as the print head ascends through various heights, ranging from 0 mm to 5 mm. These snapshots correspond to the outcomes obtained when the surface contact angle (θ_1_) is set at 90°. **(B)**. Snapshots of the simulation outcomes were obtained with a 5 mm gap (G) between the printhead and substrate. The simulation involved varying the surface contact angle (θ_1_) of the PDMS substrate, with values set at 30°, 60°, 90°, and 110°.

Here, the bioink droplet of 50 μl with a viscosity of 4 mPa·s is used with printhead diameter (D) = 6.9 mm. The selection of these parameters is tailored to the specific bioink in use and the chosen experimental conditions. The details of mDPD equations and parameterization are included in the Appendix. **Figure 2** and **Video V2** showcase the results of multiphase simulations involving a moving printhead with a wetting contact angle (θ_2_) set at 29° while varying the substrate’s wetting contact angle (θ_1_). Figure 2A(i-v) depicts snapshots from the simulation in which the liquid is drawn into the fabrication area as the print head ascends to various heights (ranging from 0 mm to 5 mm). This corresponds to the results obtained when the surface contact angle (θ1) is set at 90°. It’s important to note that this choice aligns with the contact angle range of 85°-90° observed in the different bioink formulations utilized in this study. In **Figure 2B (i-iv)**, snapshots of the results are presented, when a gap between the printhead and substrate (G) is 5 mm. The simulation was performed by varying the surface contact angle (θ_1_ = 30°, 60°, 90°, and 110°) of the PDMS substrate. Observations reveal that lower hydrophilicity (θ_1_ = 30°) promotes strong liquid adhesion with the substrate, facilitating wider droplet spreading. This necessitates a higher force to draw the liquid into the fabrication area. Conversely, higher hydrophobicity (θ_1_ = 90° and 110°) weakens liquid adhesion, making it easier to pull the bioink into the fabrication area. Higher hydrophobicity is considered better as this allows easy and smooth drawing of material into the fabrication area. However, our experiments also indicate that the upward movement of the z-stage is sufficient to pull the bioink with a contact angle (θ_1_) of 30° to the fabrication area. The mDPD model was validated using experimental data related to wetting contact angles, viscosities, and surface tensions for various bioinks. This approach eliminates the need for time-consuming trial-and-error experimental setups, thereby expediting the entire design process for 3D bioprinting experiments. Please note that the simulation results were obtained using a maximum upward movement of 5mm however during printing, stage movement of just 50 μm was necessary to print a single layer; thus for each layer, the printhead is programmed to move up by 200 μm and move down by 150 μm, and this process is repeated for each layer during the printing process.

### 3. Droplet printing of high-resolution structures with design complexities

The experimental time sequence of the structure printed from a droplet of 10% PEGDA 6K hydrogel mixed with 1 wt% of LAP (water soluble type I photoinitiator) and 0.1 wt% of tartrazine (light absorber to limit the curing depth) – a bioink formulation widely used in the field of bioprinting(**Figure 3A, Video V3**). Based on the CAD model, the droplet volume of 120 μl was pipetted in the fabrication window. (**Figure 3A(i)**). The z-stage was lowered to the initial position and the spatially patterned 405 nm laser beam with a light intensity of 2.17 mW/cm^2^ was irradiated to photo-crosslink a single layer of defined thickness. This process is repeated with synchronized upward movement of the z-stage to generate the 3D structure in a layer-by-layer fashion with an exposure time of 7s/layer. (**Figure 3A(i-iv), Video V3**). Printed structure was immersed in water for 12 hours to remove the uncrosslinked bioink and tartrazine (**Figure 3A (v)**). The bioink waste volume left behind is approximately equal to the volume needed to crosslink a single layer; with a typical layer thickness of 50μm, waste is less than 1% of initial droplet volume.

**Figure 3.**
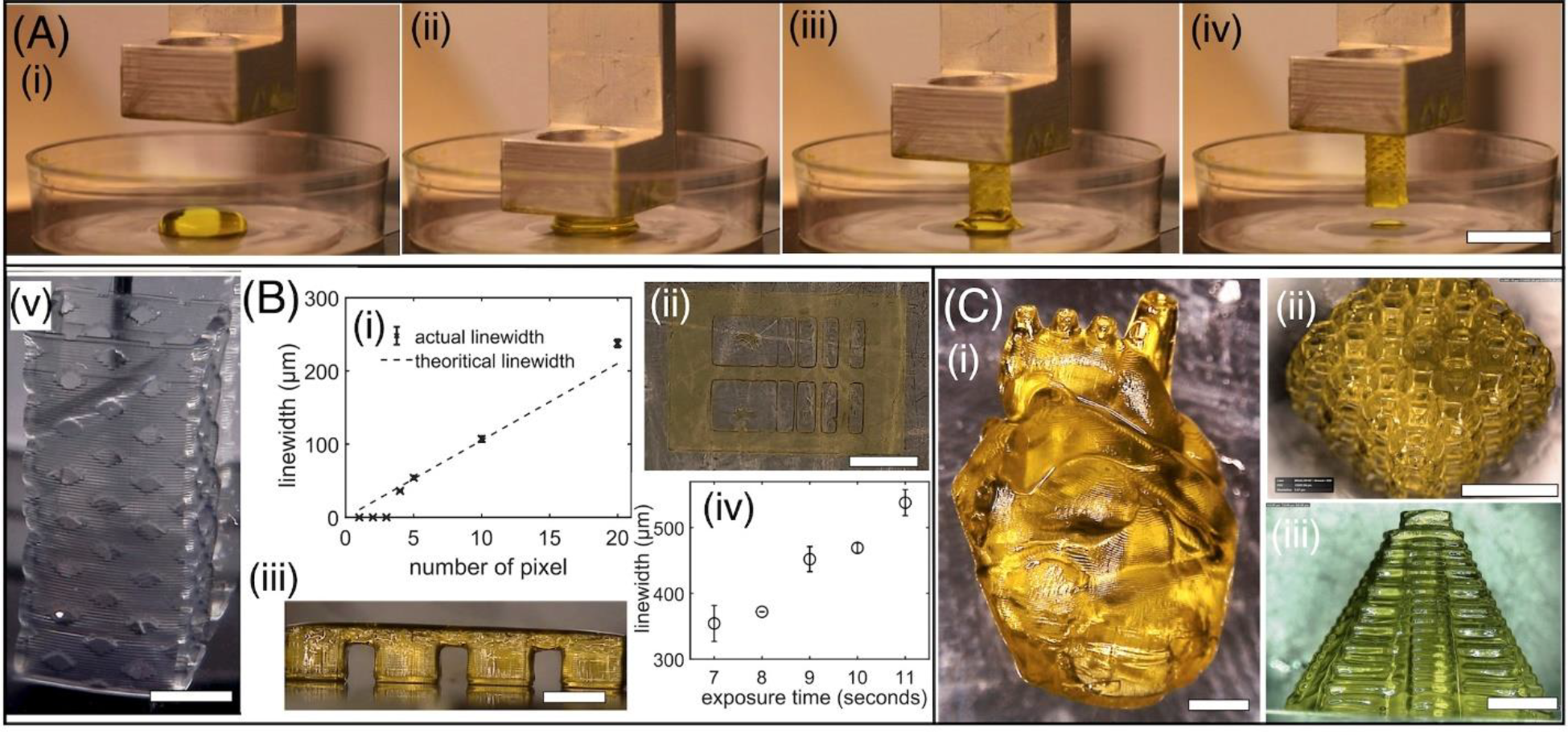
**(A)**. Sequence of images showing the droplet printing process. Scale bar - 1 cm (v) 3D printed structure after washing and removal of tartrazine light absorber. Scale bar-2 mm. **(B)** (i-ii) Plots and photo of the lateral of printed structure using PEGDA 6k MW. Scale bar-1 mm (iii-iv) Plot depicting the lateral and axial resolution of printed structure in PEGDA 400 MW. Scale bar-1.4 mm. Mean and standard deviation (n> 3). **(C)** (i-iii) Fabrication of the heart-shaped structure, lattice cube and Mayan pyramid structure in PEGDA 6k MW using DLP printing from a droplet. Scale bar-2 mm for (i), 5 mm for (ii) and 4 mm for (iii).

Printing resolution of droplet bioprinting can be influenced by multiple factors such as light exposure, stage step-size, bioink properties such as transparency, monomer reactivity, radical diffusion, and photochemical efficiency. Here, we characterize the lateral (xy) and the axial (z) resolutions using PEGDA 6k bioink formulation described above. To assess the lateral resolution, we employed a technique that involves the printing of lines ranging from 4 to 20 pixels based on a digital mask. The array of lines exhibited varying linewidths, ranging from 38±1.5 μm to 237±5.4 μm, as illustrated in **Figure 2B(i-ii)**. Results show that theoretical and printed linewidth are close to each other; this study used a laser intensity of 2.17 mW/cm^2^ and an exposure time of 7s. Notably, when attempting to achieve a feature size of 30 μm using a 3-pixel line, we encountered difficulties in preserving the integrity of the structure throughout the developmental stages. Consequently, this feature size was deemed impractical. The axial resolution, also known as the z-directional resolution, is influenced by the curing depth, which refers to the thickness of the photo-crosslinked layer. The curing depth relies on the z-directional motion of the stage, the optical absorbance of the photosensitive bioink, and the kinetics of cross-linking. It is crucial to control the curing depth to prevent unwanted crosslinking beyond the desired thickness, leading to artifacts, especially in the printing of hollow channels, undercuts and overhangs. To address this issue, we used photo-absorbers Tartrazine to increase optical absorbance and assess the curing depth using a roof-shaped structure that spans across two adjacent pillars. Due to the soft nature of PEGDA 6k bioink, it cannot maintain its mechanical integrity when printed as a single layer. (Viscosity of bioink and storage modulus of the crosslinked structure of corresponding bioinks are presented in the Method section(Table 1)) Therefore, roof structures with six layers, each with a thickness of 50 μm, were used with exposure times varying from 7-11 seconds; exposure times below 6s were not able to generate robust structures. Results show that the ideal curing thickness for the roof structure is 300 μm (6 layers; 50μm thick per layer), while the penetration depth ranged from 354±27 to 538±19 μm with increasing exposure times (7-11s) and a constant laser intensity of 2.17 mW/cm^2^ (**Figure 2B(iii-iv))**. These optimized operating conditions were used to print complex 3D structures. The structures of the human heart, lattice cube, and Mayan pyramid were printed using PEGDA 6k with 1% LAP and 0.10% tartrazine **(Figure 2C (i-iii))**. Herein, the CAD design was sliced into a layer thickness of 50 μm. Each layer was printed with the laser intensity of 2.17 mW/cm^2^ and an exposure time of 7 seconds and the structures were washed/developed by immersing the structure in a water solution. These structures demonstrate the capability of droplet printing to shape soft material into user-defined 3D structures with hollow features such as undercuts and overhangs.

**Table 1.** Viscosity, contact angle (θ_1_) of bioinks, and storage modulus of the crosslinked slab of respective bioink were used in this work. (Mean and standard deviation were calculated with n>3 independent experiments. *Sample size of n=1 due to drying at the edge. Out of five samples only one was successfully completed.)

| Bioink | Associated Figure | Viscosity (mPa*s) | Contact angle (°) | Storage Modulus of the crosslinked slab at 1 Hz (KPa) |
| --- | --- | --- | --- | --- |
| 10% PEGDA 6K | Figure 3 | 3.19±0.06 | 85±3.15 | 3.11±0.18 |
| 5% PEGDA6k+5%GELMA (1:1) | Figure 5, Figure 7 | 2.11±0.044 | 91.23±0.44 | 0.24±0.019 |
| 7.5% GelMA, 10% PEGDA 6K, and 20% PEGDA 700 (6:3:1) | Figure 6 | 7.8929* | 83.29±0.31 | 2.70±0.122 |

### 4. Optimization of formulation and printing conditions for cell-laden bioinks

Due to use of small-volume droplets, bioink optimization becomes a rapid, low-cost, and simple process. Here, we used model cell line (C3H/10T1/2) to screen a range of bioink formulations using varying amounts of PEGDA 6K hydrogel, GelMA hydrogel, and LAP photoinitiator; GelMA provides cell-adhesive matrix for encapsulated cells, while PEGDA 6k provide high print fidelity. We optimized exposure time, hydrogel composition, and photoinitiator concentration using model C3H/10T1/2s at an initial concentration of 1 × 10^6^ cells/ml (before mixing with bioink); cell solution: bioink ratio was 2:8. <u>Optimization of Exposure dose.</u> Rectangular slabs of 1 mm height (20 layers with a layer thickness of 50 μm) were printed using droplet bioprinting by varying the exposure time from 2.5 to 4 seconds while maintaining a constant laser intensity of 2.17 mW/cm^2^ **(Figure 4A)**. Structures printed using the exposure time of 2.5 seconds exhibit high cell viability. However, the structure was partially crosslinked, resulting in excessive swelling and deformation upon immersion in the media solution. The longer exposure time of 4 seconds resulted in a stiff structure with poor cell viability. An exposure time of 3 seconds per layer was found to be ideal in terms of mechanical integrity and cell viability **(Figure 4A)**. <u>Optimization of bioink concentration</u>. A similar structure was printed by varying the concentration of the hydrogel (3, 5, 7.5 and 10%: GelMA + PEGDA) while maintaining constant exposure time of 3 seconds and laser intensity of 2.17 mW/cm^2^. Bioink formulation of 3% GelMA and 3% PEGDA was too soft to handle while bioink formulation of 7.5% GelMA and 7.5% PEGDA showed decreased cell viability. Bioink formulation with 5% GelMA and 5% PEGDA was chosen for subsequent studies based on their cell viability and printing fidelity. **(Figure 4B)**. <u>Optimization of LAP concentration.</u> Here, we varied the concentration of LAP (0.25%, 0.5% and 1% LAP) while bioink (5% GelMA + 5% PEGDA), laser intensity (2.17 mW/cm^2^) and exposure time (3s/layer) was held constant. **(Figure 4C)**. Based on the results, a LAP concentration of 0.5% was chosen for next set of experiments.

**Figure 4.**
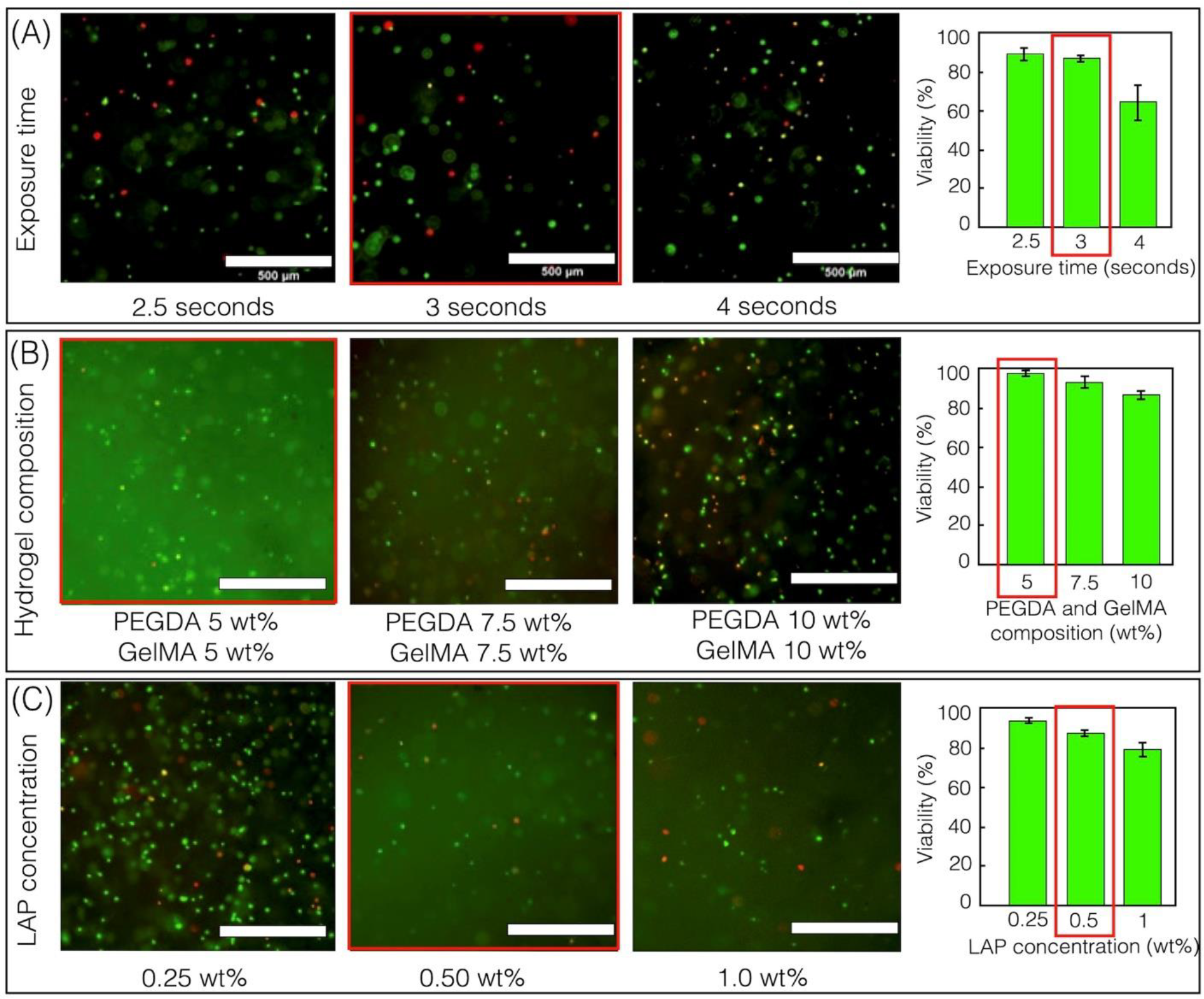
Optimization of (A) laser intensity and exposure time. (B) Bioink composition. and (C) photoinitiator concentration. Optimized parameters are marked with red rectangles. Scale bars-500 μm. (Mean and standard deviation were calculated with n>3 independent experiments.)

### 5. Droplet bioprinting of C3H/10T1/2 – laden 3D constructs

Using optimized conditions, droplet bioprinting was used to print a cell-laden 3D structure with spatially patterned channels. Prior to bioprinting, the sample holder and PDMS slab were sterilized under UV radiation. Then, a 130μl volume of bioink, maintaining a C3H10T1/2: bioink ratio of 2:8 (1x10^6^ cells/ml before mixing) was pipetted in the fabrication window, and a rectangular slab structure of height 1 mm (20 layers with a layer thickness of 50 μm) with multiple wells spaced apart by 1mm and 300μm were printed using a laser intensity of 2.17 mW/cm^2^, an exposure time of 3s/layer, and a layer thickness of 50 μm. (**Figure 5A, B)**. After printing, the structures were washed with PBS for 5 minutes, cultured for 7 days, and imaged using confocal microscopy. Results show a uniform distribution of encapsulated cells throughout the structure (**Figure 5C**). In addition, cell viability was assessed at different spatial locations marked by red (dotted) and blue (solid) rectangles at a depth of 500 μm (**Figure 5D**). Cell viability of the spatial location marked by the red rectangle increased from 51.66±2.25% to 91±1,69% from Day 1 to 7 respectively. **(Figure 5D (ii-v)**. Encapsulated cells were found to be more viable with dendritic morphologies in areas marked by the blue rectangle (96±1.24%), as the wells were closer together with greater access to media, (**Figure 5D (iv,v)**).

**Figure 5.**
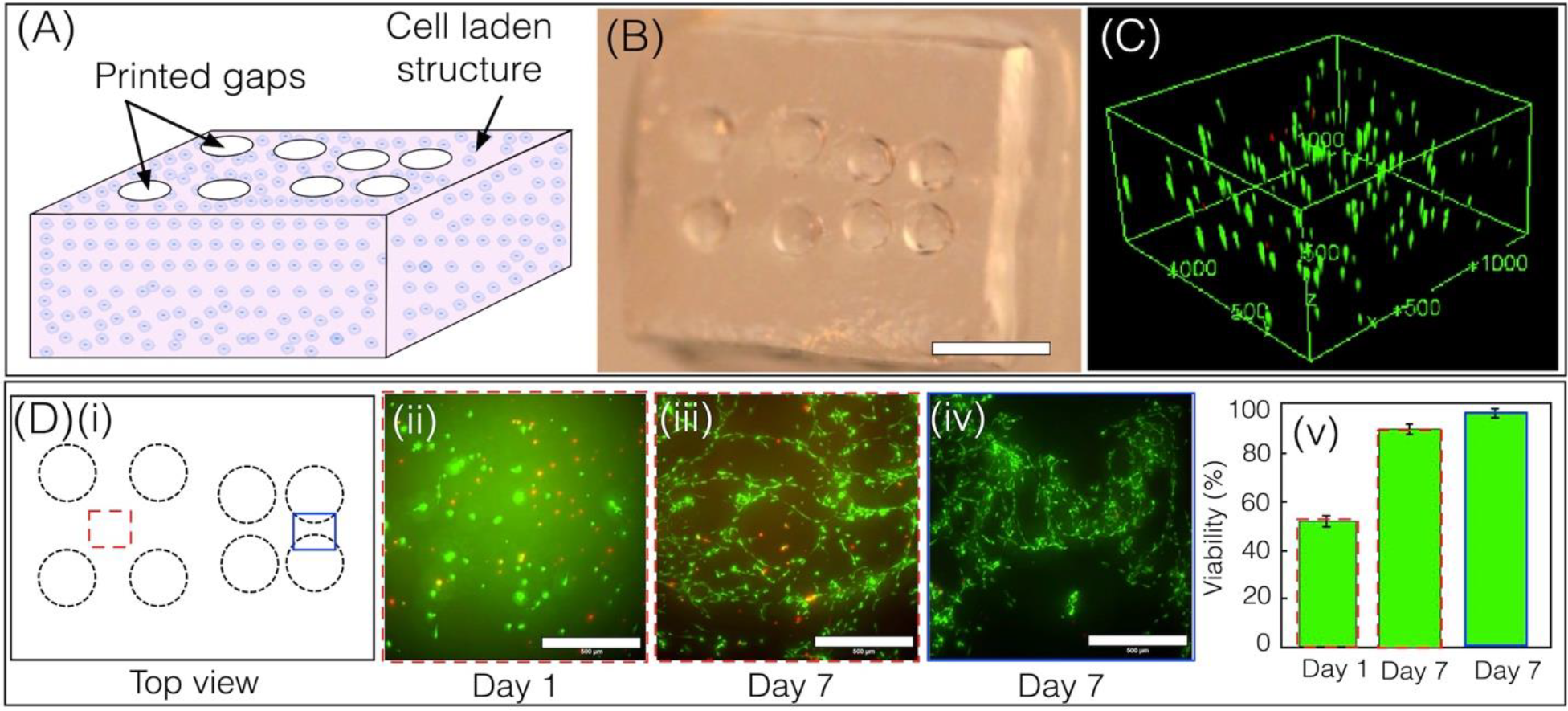
**(A**,**B**) Illustration and photo of the 3D printed construct made with a GelMA/PEGDA mixture and 10T1/2 cell solution (scale bar-2.5 mm). **(C)** Confocal reconstructed images showing uniform distribution of cells within the 3D printed construct. Units are shown in micrometers. **(D)(i)** Top view cartoon of the bioprinted structure. **(ii-v)** Live-Dead stained images showing the viability of encapsulated cells on Days 1 and 7 at different locations, with comparative viability plots.(Scale bars - 500 μm). (Mean and standard deviation were calculated with n>3 independent experiments.)

### 6. Microfluidic chips with endothelialized channel networks

In this work, we used droplet bioprinting to print 3D chips that consist of embedded microchannel networks, a structure widely used in vascular tissue engineering. (**Figure 6A**) The bioink formulation consists of 7.5% GelMA, 10% PEGDA 6K, and 20% PEGDA 700 (6:3:1). PEGDA 700 was added to provide additional structural strength and increase printing fidelity. The structure was printed using 150 μl of the bioink with a laser intensity of 2.17 mW/cm^2^, layer thickness of 50 μm and exposure time of 2.5 seconds. To emphasize the channels within the structure, a red food color dye was infused **(Figure 6B)**. Following printing, the structure underwent overnight sterilization using UV light. To promote cell adhesion around the channel, rat tail collagen was mixed to a concentration of 0.5 mg/ml in a 20 mM acetic acid solution and carefully pipetted into the channels. After 30 minutes of incubation, the channels were fully aspirated, and a thorough rinse with PBS was performed. Following that, endothelial cell solution (2H11s, cell density-10 M/ml) was perfused into the channels and allowed to incubate for 20 minutes to facilitate cell adhesion. The structure was then inverted and allowed to incubate for an additional 20 minutes. The cell solution was subsequently aspirated, and fresh medium was introduced into the channels. The structure was cultured under standard conditions for 5 days with medium changes every 2 days. Further, the structure was inverted every two days to achieve a homogeneous distribution of endothelial cells along the channel wall. On Day 5, the cells were fixed, stained for actin and nuclei, and confocal imaging was used to characterize endothelial morphology within the channel network. Results show that the endothelial monolayer covers the entire channel surfaces **(Figure 6C, D, Video V4)**.

**Figure 6.**
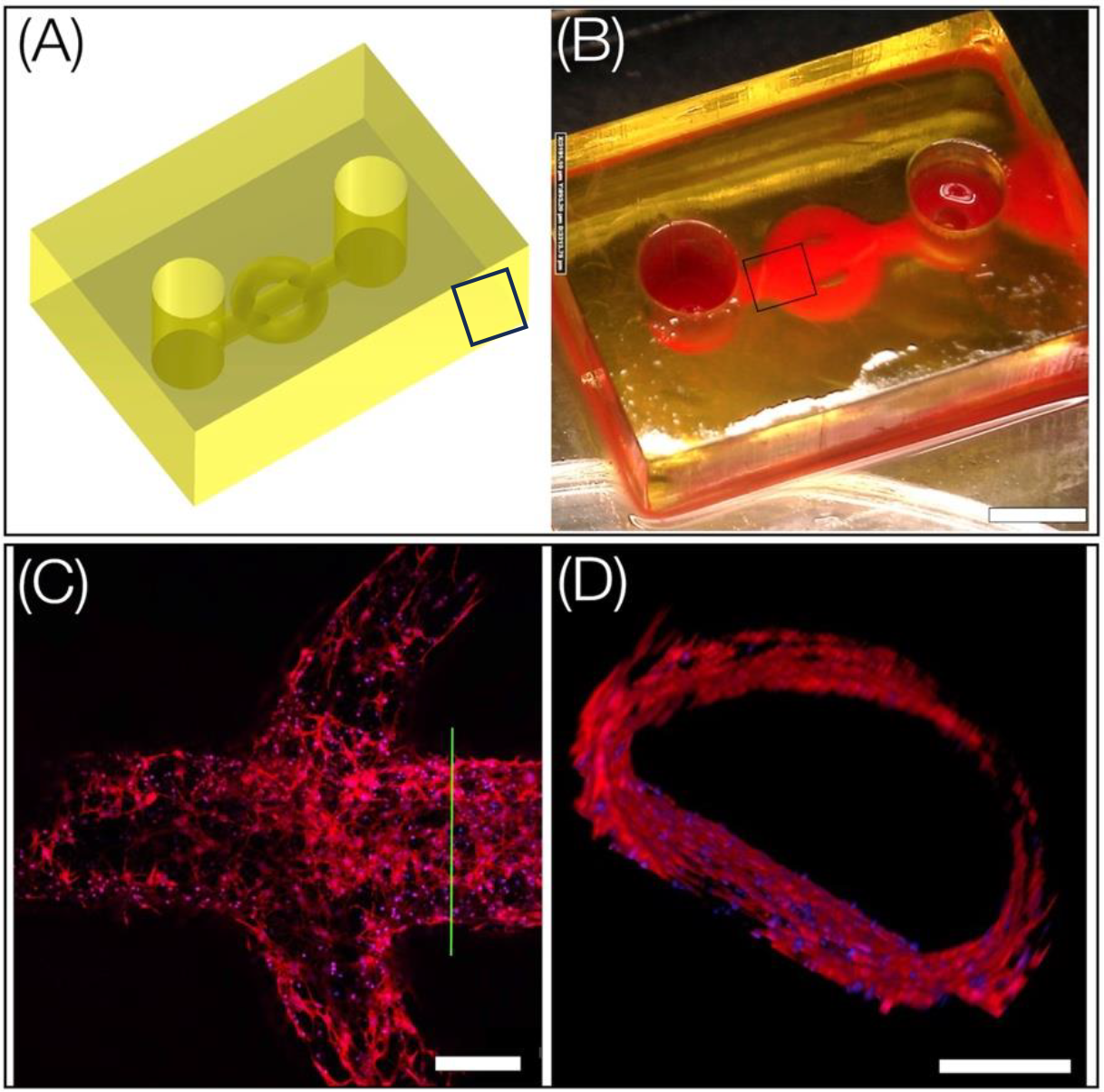
**(A-B)** CAD and printed chip with embedded channel network. Red food color dye was used to visualize the flow path (Scale bar -2 mm). **(C)** Confocal image of a section of the 2H11 lining from a single plane (scale bar-100 μm). **(D)** Cross section at green line showing the formation of lumens by 2H11 endothelial cells around the fabricated channel (Red-actin and Blue-nuclei) (Scale bar - 200 μm).

### 7. Droplet bioprinted structures subjected to osteogenic induction exhibit expected mineral deposition

To demonstrate that bioprinted structures with relevant functional outcomes can be generated, we printed a rectangular slab using bioink laden with model osteoblasts (human osteosarcoma, Saos-2, ATCC) **(Figure 7A, B)**. First, the printing process was rapidly optimized using a low-volume bioink (80 μl), using a laser power of 2.17 mW/cm^2^ and an exposure time of 3s/layer. A total of 20 layers were printed with a layer thickness of 50 μm. Post-printing, structures were cultured in normal growth medium for 48 hours prior to introducing them to well plates with osteogenic media for a duration of three weeks. Results exhibit high viability of encapsulated Saos-2 cells with mineral deposition throughout the structure, as assessed by brightfield images, SEM-EDS analysis, and micro-CT **(Figure 7)**. In contrast to control acellular samples with no mineral, printed construct, the EDS spectra, and elemental maps revealed the presence of calcium and phosphorus minerals deposited by encapsulated Saos-2. **(Figure 7 F (ii-iv), Figure S1)**. Calcium-phosphorus crystal nucleation is known to begin within osteogenic cells and later migrate to the extracellular matrix resulting in the formation of hydroxyapatite crystals.^[21]^ Micro-CT also confirmed the presence of mineral formation **(Figure 7G, Video V5)**.

**Figure 7.**
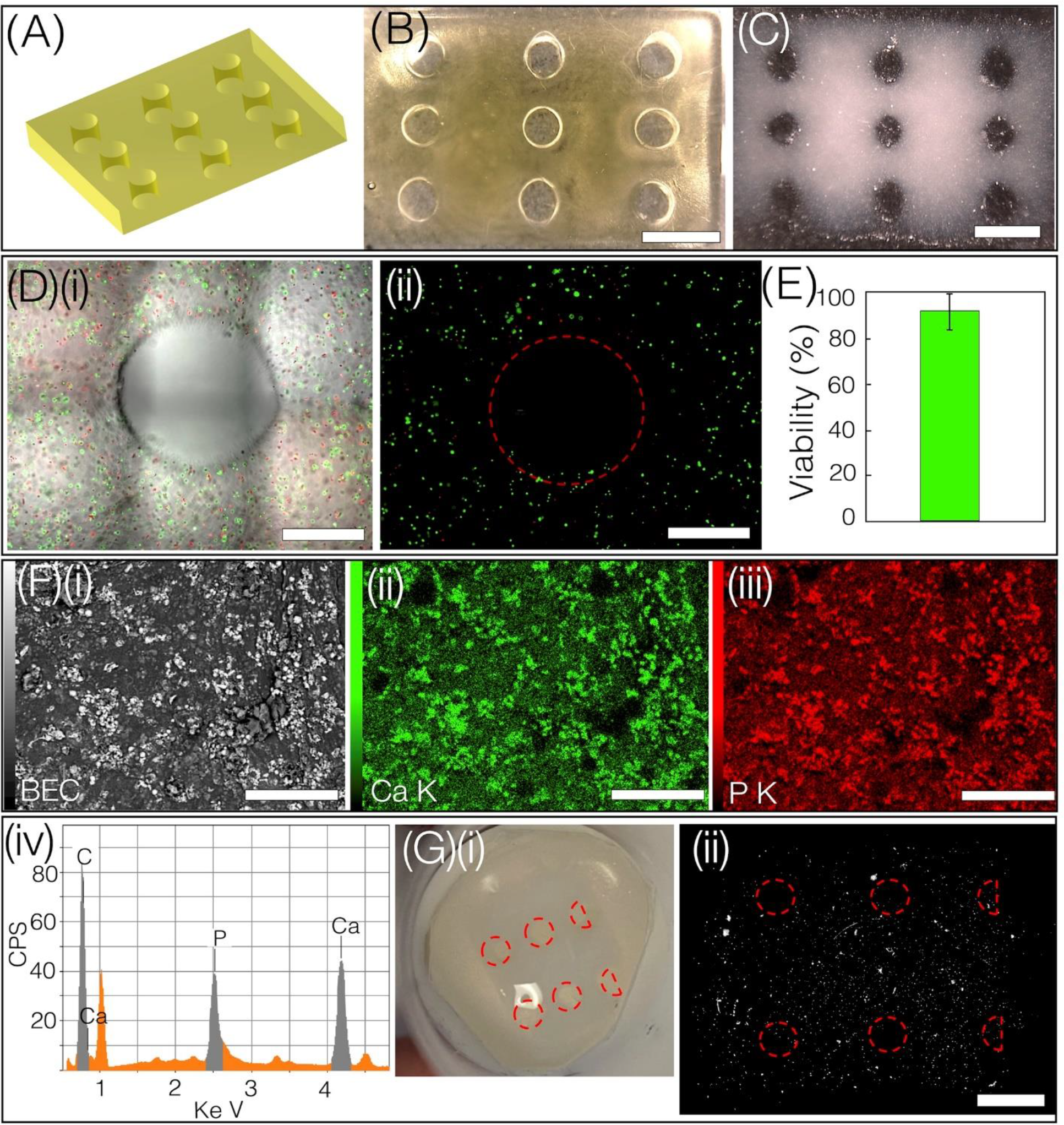
(A-B) CAD and droplet bioprinted construct, scale bar-2 mm. (C) The brightfield image of the construct after 21 days of osteogenic static culture, depicting opacity due to mineral deposition. (Scale bar-2mm) (D) (i-ii) Merged and fluorescent image of one channel. (Scale bar-500μm). (E) Cell viability within the printed construct on Day 2 post-printing. (F)(i) A scanning electron microscope (SEM) image was used to image the structure (a small section) shown in Figure (C), followed by (ii) calcium map (iii) potassium map, (Scale bar-300μm) and (iv) EDS spectrum depicting the presence of Calcium and Phosphorus. (G)(i) Photographs of mineralized structure. (ii) Micro-Ct image of mineralized structure (Scale bar-2 mm).

## CONCLUSION

While extrusion bioprinters remain the most widely used in the field,^[22–25]^ many vat-based photopolymerization methods have emerged due to their ability to print high-resolution and complex design architectures. However, current vat-based bioprinters face many challenges such as low fidelity printing with soft bioinks, low cell viability, large waste and high costs due to the requirement of filling the vat with expensive bioinks even for small size prints, print defects due to gravity-based cell settling and/or changes in bioink properties during printing.^[11,14,23,26]^ Here we report a vat-free, droplet bioprinting method that can print acellular and cell-laden constructs with minimal waste. The up-and-down motion during printing allows for efficient mixing of the cell-laden inks to generate constructs with uniform cell density with over 90% viability. A new model was developed to simulate the dynamic process of droplet bioprinting. Overall, this low-volume, low-cost method will be ideal for screening bioink formulations with a large number of design variables, as well as making customized, high-resolution 3D constructs for a range of biomedical applications.

## Methods

### 1. GelMA synthesis

Gelatin Methacrylate (GelMA) macromer was synthesized using a previously reported protocol.^[27]^ Briefly, 10 g porcine skin gelatin (Sigma Aldrich, St. Louis, MO) was mixed in 200 ml phosphate buffered saline (PBS, Thermo Fisher Scientific), stirred at 45?, and methacrylic anhydride was added to the solution, and stirred for 3 h. After stirring, the mixture was dialyzed against distilled water for 1 week at 40? to remove the unreacted groups from the solution. The dialyzed GelMA was lyophilized in a freeze dryer (Labconco, Kansas City, MO) for one week. To prepare 15% (w/v) GelMA, a stock solution was prepared by mixing 1.5 g freeze-dried GelMA with 10 ml of deionized water (dissolved at 40?), and 0.25% (w/v) UV photoinitiator lithium phenyl-2,4,6-trimethyl-benzoyl phosphinate (LAP) was added into the solution. GelMA pre-polymer solution was diluted using DI water to obtain either 7% or 10% GelMA and filtered (pore size=0.2 μm) and used within 2 h after preparation.

### 2. LAP synthesis

Lithium phenyl-2,4,6-trimethyl-benzoyl phosphinate (LAP) was synthesized in a two-step process according to the literature.^[28]^ At room temperature and under nitrogen, 2,4,6-trimethyl benzoyl chloride (4.5 g, 25 mmol) was added dropwise to continuously stirred dimethyl phenyl phosphonate (4.2 g, 25 mmol). The reaction mixture was then stirred for 24 hours therefore an excess of lithium bromide (2.4 g, 28 mmol) in 50 mL of 2-butanone was added to the reaction mixture from the previous step which was then heated to 50 ?. After 10 min, a solid precipitate formed. The mixture was then cooled to room temperature, and then filtered. The filtrate was washed with 2-butanone (3 x 25 mL) to remove unreacted lithium bromide and dried under vacuum to give LAP (6.2 g, 22 mmol, 88%) as a white solid.

### 3. PEGDA 6K synthesis

PEGDA (6000 MW) was synthesized in the laboratory and subsequently dissolved in water to attain a 10% w/w concentration of PEGDA 6k, along with 1% w/w LAP.^[29]^ To synthesize PEGDA 6k, 24 grams of PEG 6k were dissolved in 100 milliliters of anhydrous DCM and cooled in an ice bath. Under a nitrogen blanket and with continuous stirring, 1.12 milliliters of triethylamine were introduced into the chilled solution, followed by the gradual addition of a solution containing 1.27 milliliters of acryloyl chloride, which had been diluted to a final volume of 15 milliliters in anhydrous DCM via an addition funnel. The reaction mixture was allowed to stir for 30 minutes under a nitrogen blanket, after which it was left to proceed overnight with continuous stirring in a covered flask under nitrogen. Following this, quaternary ammonium salts were removed from the reaction mixture using liquid-liquid extraction into 16 mL 2 M K_2_CO_3_. Once the emulsion separated, the organic PEGDA-containing layer was isolated, dried with anhydrous Na_2_SO_4_, and then filtered. PEGDA was then precipitated through dropwise addition to hexane with vigorous stirring. The resulting precipitate was separated via vacuum filtration, followed by washing with three aliquots of hexane and then three aliquots of diethyl ether at -80°C. Finally, the washed PEGDA was dried under vacuum conditions at room temperature for one week.

### 4. Relevant properties of bioinks

### 5. Optical setup

The bioprinter employed in this study utilizes a 405 nm laser source provided by Toptica, capable of generating a continuous-wave (CW) laser beam with a maximum power output of 300 mW. To achieve the desired laser beam characteristics, several components are integrated into the system. Firstly, a shutter (SH05, Thorlabs) is strategically positioned after the laser source to collimate and expand the beam. To further refine the beam profile, a 2f-transfer lens assembly, consisting of lenses with focal lengths of 40 mm and 200 mm, is utilized. To achieve a uniform intensity distribution from the initial Gaussian laser beam, spatial filtering is implemented using a 25 μm pinhole. Additionally, to mitigate the occurrence of laser speckles, a diffuser (provided by RPC Photonics Inc.) is employed. The diffuser is mounted on a rotating mount, which helps minimize the speckle effect. The modified laser beam is directed toward a Digital Micromirror Device (DMD) manufactured by DLi Inc. This DMD, known as the 0.95’’ 1080p UV DMD, consists of an array of micromirrors capable of spatially patterning laser beams. Following the patterning process, the laser beam is projected onto infinity-corrected projection optics, which consist of two lens systems with focal lengths of 300 mm each. These lens systems are positioned at a distance of 60 cm from each other. To precisely focus the spatially patterned laser beam within the fabrication window, the distance between the lenses is meticulously adjusted. The fabrication process involves a PDMS slab positioned above a heater (WP-16 Warner instrument), which is heated to a temperature of 40ºC specifically for GelMA printing. This is because GelMA typically undergoes gelation at room temperature. The movement and coordination of the PDMS slab are facilitated by an L-shaped stage, which is controlled by a three-dimensional linear stage provided by PI. The stage’s movements are coordinated using a custom LabVIEW code, ensuring precise positioning and alignment.

### 6. Rheological Testing

Rheological characterization was performed using a TA Instruments Discovery Hybrid Rheometer (DHR-3) with a temperature-controlled lower Peltier plate (TA Instruments, New Castle, DE, USA). Resin viscosities were measured using 40 mm parallel plate geometry and a flow sweep from a shear rate of 0.1 to 100 s^-1^. All viscosity measurements were performed at 25°C except for resin formulations containing GelMA, which were instead performed at 37°C to prevent physical gelation. To obtain storage modulus, frequency sweep experiments were performed on photo-crosslinked hydrogel samples using 8 mm parallel plate geometry at 25°C in the range of 0.1 to 100 Hz. Cylindrical photo-crosslinked hydrogel samples were prepared via exposure to 405 nm light patterned using an 8 mm diameter circular mask in a 1 mm deep PDMS well. The resulting hydrogel samples were 8 mm in diameter and 1 mm thick. Unless otherwise noted, all rheological measurements were performed in triplicate.

### 7. Micro-CT analysis

The cellularized construct was removed from the static media dish intact, fixed in formaldehyde (4% for 24hr), washed in PBS, and placed on a solid 3D-printed circular base. The base was placed inside a 20 mm diameter sample holder for micro-CT imaging (micro-CT 40, Scanco Medical AG, Brüttisellen, Switzerland) and kept hydrated with PBS, utilizing a foam spacer positioned on top to prevent sample movement. Samples were imaged at a 10 μm isotropic voxel resolution (55 kV, 145 mA, 200 ms integration time). After scanning, the reconstructed micro-CT images (.isq files) were imported into Materialise Mimics, a 3D medical image segmentation software, for analysis.

The images were cropped to isolate the construct, and a lower global threshold of 200 mg HA cm-3 was applied to identify minerals within. A 3D reconstruction was created from this data and exported as a .stl file for visualization.

### 8. EDS analysis

To perform EDS analysis, mineralized cell-laden constructs were subjected to solvent exchange via serial incubation in ethanol solutions of increasing concentration (10%-100% v/v ethanol in MilliPore water in 10% increments) for 30 minutes each. Solvent exchanged samples were dried in vacuo for 24 hours, then mounted on aluminum stubs with carbon tape and sputter coated with Au/Pd (Denton Vacuum Desk V, 25mA; 60s). Coated samples were examined on a JEOL JSM-IT100LA scanning electron microscope under high vacuum at 20 kV using compositional backscatter imaging (BEC) for increased contrast between mineralized and non-mineralized regions. Energy-dispersive X-ray spectroscopy (EDS) was used to detect and map calcium, phosphorous, and carbon within the samples. Cell-free control samples were analyzed under the same conditions.

### 9. Cell Culture and Staining

#### C3H/10T1/2 cell line

*C3H/10T1/2*s were maintained using a cell growth media consisting of Basal Medium Eagle (BME, Gibco™) supplemented with 10% heat-inactivated fetal bovine serum (FBS, Atlanta Biologicals), 1% Glutamax (Gibco™) and 1% penicillin-streptomycin (10,000 U/mL, Gibco™). The cells were maintained in a controlled environment at 37°C with 5% CO_2_ and 90 % humidity. The cells were harvested before reaching full confluency with 0.05 % trypsin.

#### Saos-cell line

We utilized Saos-2 as a suitable model cell line for osteogenesis. The cultivation of Saos-2 cells involved using Dulbecco’s modification of Eagle’s medium (DMEM, Gibco™) as the foundational medium, supplemented with 10% heat-inactivated fetal bovine serum (FBS, Atlanta Biologicals), 1% Glutamax (Gibco™) and 1% penicillin-streptomycin (10,000 U/mL, Gibco™). The cells were maintained in a controlled environment at 37°C with 5% CO_2_ and 90 % humidity. The cells were harvested using 0.25% trypsin for new experiments. To induce chemical mineral production in Saos-2 cells, we introduced specific supplements to the base medium. These supplements included 100 μM L-ascorbic acid-2-phosphate (AA2P, Sigma-Aldrich), 5 mM β-glycerophosphate (BGP, Sigma-Aldrich). ***2H-11 cell line***. We cultured and harvested the 2H-11 cell line using the same method and base media employed for the Saos cell line.

#### Fluorescence staining

To study cellular morphology, the cells were stained for F-actin and nuclei. The cells were first fixed with 4% formaldehyde for 30 minutes and then treated with 0.2% Triton X-100 in DPBS for 30 minutes to permeabilize the cells. Subsequently, the cells were stained with either phalloidin (Alexa-Fluor 568, Invitrogen) at a dilution of 1/100 or with phalloidin-rhodamine at a dilution of 1/250 in 10% horse serum for 45 minutes at room temperature to visualize f-actin, and DAPI (Life Technologies) at a dilution of 1/1000 for 5 minutes at room temperature to visualize cell nuclei. The images of the fluorescently stained sample were acquired using a confocal microscope (Zeiss LSM 980).

#### Live/Dead analysis

A Live/Dead assay kit from Invitrogen, which includes Calcein AM and Ethidium homodimer, was employed to assess the cell viability. To summarize, a solution containing 1/2000 of Calcein AM and 1/1000 of Ethidium homodimer in phenol red free cell media was carefully added to the cell-laden structure. The structure was then incubated at 37°C for 45 minutes, after which images were acquired using an inverted microscope (Nikon Eclipse).

## Acknowledgment

We are thankful to the National Institutes of Health (R21 GM141573-01) and the Syracuse University Collaboration of Unprecedented Success and Excellence (CUSE) and the BioInspired Seed grant program for the financial support for this project.

