## Supplementary data for "Droplet bioprinting of acellular and cell-laden structures at high-resolutions"

### 1. EDS analysis of mineralized structure as compared to a control sample

(A) Mineralized Sample

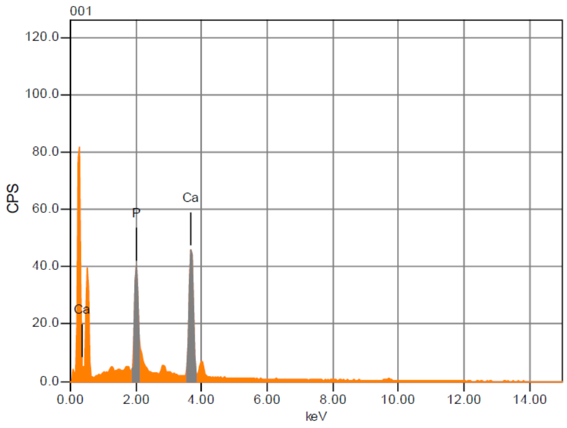


| Formula | mass% | Atom% | Sigma |
|---|---|---|---|
| C* | 76.38 | 90.78 | 0.02 |
| P | 7.67 | 3.53 | 0.02 |
| Ca | 15.95 | 5.68 | 0.03 |
| Total | 100.00 | 100.00 | |

(B) Control Sample

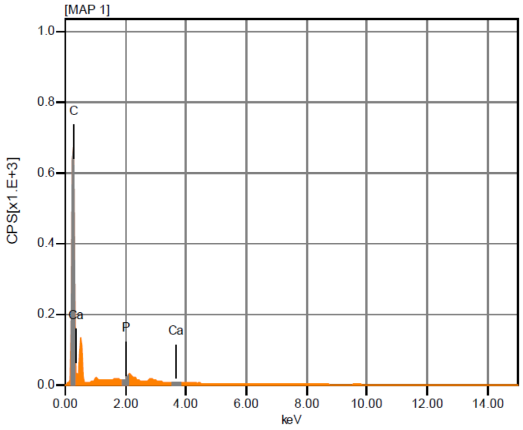


| Formula | mass% | Atom% | Sigma |
|---|---|---|---|
| C* | 99.92 | 99.98 | 0.01 |
| P | nd | nd | |
| Ca* | 0.08 | 0.02 | 0.00 |
| Total | 100.00 | 100.00 | |

***Figure S1. (A)** EDS spectrum from the as-fabricated mineralized structure shows the presence of Calcium and Phosphorus peak. **(B)** EDS spectrum from the control structure (cell free) shows the presence of Calcium and Phosphorus peak.*

**Appendix: Many-Body Dissipative Particle Dynamics**

Many-body dissipative particle dynamics (mDPD) method is developed based on the standard DPD framework [1]. The time evolution of mDPD particles $i$ is governed by the conservation of momentum, which is described by the following set of equations,

$$\frac{d\boldsymbol{r}_i}{dt} = \boldsymbol{v}_i \tag{1}$$

$$\frac{d\boldsymbol{v}_i}{dt} = \boldsymbol{F}_i = \sum_{i\neq j}\left(\boldsymbol{F}_{ij}^C + \boldsymbol{F}_{ij}^D + \boldsymbol{F}_{ij}^R\right) \tag{2}$$

where $t$, $\boldsymbol{r}_i$, $\boldsymbol{v}_i$and $\boldsymbol{F}_i$ denote time, and position, velocity, force vectors, respectively. The summation of forces is carried out over all other particles within a cutoff radius $r_c$, beyond which the direct interactions between particles are considered to be zero.

The three components of $\boldsymbol{F}_i$ include the conservative force $\boldsymbol{F}_{ij}^C$, dissipative force $\boldsymbol{F}_{ij}^D$ and random force $\boldsymbol{F}_{ij}^R$, which are expressed as [2]

$$\boldsymbol{F}_{ij}^C = A\omega_c(r_{ij})\boldsymbol{e}_{ij} + B(\rho_i + \rho_j)\omega_d(r_{ij})\boldsymbol{e}_{ij} \tag{3}$$

$$\boldsymbol{F}_{ij}^D = -\gamma\omega_D(r_{ij})(\boldsymbol{e}_{ij}\cdot\boldsymbol{v}_{ij})\boldsymbol{e}_{ij} \tag{4}$$

$$\boldsymbol{F}_{ij}^R = \delta\omega_R(r_{ij})\xi_{ij}\Delta t^{-\frac{1}{2}}\boldsymbol{e}_{ij} \tag{5}$$

where $r_{ij}$ is distance between particles $i$ and $j$, $\boldsymbol{e}_{ij}$ is the unit vector from particle $j$ to $i$, and $\boldsymbol{v}_{ij} = \boldsymbol{v}_i - \boldsymbol{v}_j$ is the velocity difference. $\omega_c$, $\omega_d$, $\omega_D$ and $\omega_R$ are the weight functions of $\boldsymbol{F}_{ij}^C$, $\boldsymbol{F}_{ij}^D$and $\boldsymbol{F}_{ij}^R$, respectively. A negative coefficient $A < 0$ stands for an attractive force and a positive coefficient $B > 0$ results in a density-dependent repulsive force. $\gamma$ is dissipative parameter, $\delta$ is the Gaussian white noise with zero mean and unit variance which describes the degrees of freedom that have been eliminated from the coarse-graining process [3]. The dissipative force and random force act as a thermostat if the dissipation parameter $\gamma$ and the amplitudes of white noise $\delta$ satisfy the fluctuation-dissipation theorem [4] requiring $\delta^2 = 2\gamma k_B T$ and $\omega_D(r) = [\omega_R(r)]^2$, in which $k_B$ is the Boltzmann constant and $T$ is the temperature. A common choice of the weight function is $\omega_c = 1 - r_{ij}/r_c$ and $\omega_D = {\omega_R}^2 = \left(1 - r_{ij}/r_c\right)^s$for $r_{ij} \leq r_c$ and vanish for $r_{ij} > r_c$, while $\omega_d = 1 - r_{ij}/r_d$ with a different cutoff radius $r_d < r_c$.

The mDPD system a droplet-based 3D printing model is constructed with parameters $A_{ss} = A_{ll} = -40, B = 25, \gamma = 18, \delta = 6, k_B T = 1, r_c = 1.0, r_d = 0.75, s = 0.5,$ and $A_{sl}$ changes from $-22$ to $-34$ to adjust the wettability of solid surfaces, in which the parameters $A_{ss}$, $A_{ll}$ and $A_{sl}$ represent the attractive coefficients in Eq. (3) for solid-solid, liquid-liquid and solid-liquid interactions. Varying the strength of attractive interactions between solid and liquid particles can tune the solid-liquid interfacial tension and hence change the wetting contact angles. The mDPD system with an initial meniscus volume of 50 microliters consists of 793,746 particles, in which 169,169 particles form the cross-linked printhead, 198,888 particles being the PDMS substrate, and 425,689 particles representing the liquid meniscus. The parameters in the mDPD model are calibrated based on the fluid properties of bioink with a viscosity of $4\ \mathrm{mPa} \cdot \mathrm{s}$, and the wetting contact angle $\theta_1$varies from $30°$ to $110°$ for different PDMS base while $\theta_2$ is $29°$ for crosslinked slab.
